# Host defense triggers rapid adaptive radiation in experimentally evolving parasites

**DOI:** 10.1101/420380

**Authors:** Sarah E. Bush, Scott M. Villa, Juan C. Altuna, Kevin P. Johnson, Michael D. Shapiro, Dale H. Clayton

## Abstract

Adaptive radiation occurs when the members of a single lineage evolve different adaptive forms in response to selection imposed by competitors or predators. Iconic examples include Darwin’s finches, Caribbean anoles, and Hawaiian silverswords, all of which live on islands. Parasites, which live on host “islands,” show macroevolutionary patterns consistent with adaptive radiation in response to host-imposed selection. Here we show rapid adaptive divergence of experimentally evolving feather lice in response to preening, the main host defense. We demonstrate that host defense exerts strong phenotypic selection for crypsis in lice transferred to different colored rock pigeons (*Columba livia*). During four years of experimental evolution (∼60 generations), the lice evolved heritable differences in color. The color differences spanned the phenotypic distribution of congeneric species of lice adapted to other species of pigeons. Our results indicate that host-mediated selection triggers rapid divergence in the adaptive radiation of parasites, which are among the most diverse organisms on earth. Our research suggests that host defense should be included with competition and predation as a major mechanism driving the evolution of biodiversity by adaptive radiation.

## MAIN TEXT

Adaptive radiation is a major source of organismal diversity [1-5]. Ironically, however, the role of this process in parasite diversification remains unclear, despite the fact that parasites are among the most diverse organisms on Earth [6-10]. Parasites may adapt and radiate among host species, just as free-living species adapt and radiate among islands within archipelagos. Host species are analogous to islands that limit dispersal and gene flow between parasite populations and species. However, as in the case of physical islands, the barriers created by host islands are not absolute because even host-specific parasites occasionally switch host lineages over macroevolutionary time [11-18]. Host switching can lead to patterns consistent with adaptive radiation in phytophagous insects [15, 16, 19], fungal plant pathogens [13], helminth worms [18], and ectoparasitic arthropods [11, 14]. Host-mediated selection also appears to drive the adaptive divergence of parasites exposed to varying defensive regimes on different host islands [6, 9, 17, 20]. This hypothesis has not been tested experimentally because it is difficult to isolate and manipulate components of host defense. We conducted such a test using an unusually tractable host-parasite system consisting of rock pigeons (*Columba livia*) and their feather lice (Insecta: Phthiraptera: Ischnocera).

Feather lice are host-specific parasites of birds that feed on the downy regions of feathers, causing energetic stress that leads to a reduction in host fitness through reduced survival and mating success [11]. Feather lice depend on feathers for efficient locomotion. Thus, transmission between host individuals usually requires direct contact, such as that between parent birds and their offspring in the nest. However, feather lice can also disperse by hitchhiking phoretically on parasitic flies that are less host-specific than lice [21]. As a consequence, lice periodically end up on novel host species [11]. Birds combat lice by removing them with their beaks during regular bouts of preening. Lice are thought to escape from preening through background matching crypsis because light colored bird species have light colored lice, whereas dark colored species have dark colored lice [22] (Fig.1A,B). Although these observations suggest that preening is the selective agent responsible for the evolution of cryptic coloration in feather lice, this hypothesis has never been tested experimentally.

We measured the selective effect of preening on the color of pigeon lice (*Columbicola columbae*) by comparing the survival of experimentally manipulated lice placed on different colored rock pigeons (see Materials and Methods). Live lice were painted black or white and distributed evenly among 8 black and 8 white pigeons (Fig. 2A-D). Half of the birds could preen normally, whereas the other half had their preening ability impaired with harmless bits that prevent complete closure of the beak (Fig. 2E,F). After 48 hours, all birds had their lice removed by body washing [23]. Birds with normal preening had significantly more cryptic lice than conspicuous lice at the end of the experiment (Fig. 2G). Conspicuous lice were 40% more likely be removed by preening, revealing intense selection for cryptic coloration. In contrast, there was no significant difference in the number of cryptic and conspicuous lice on pigeons with impaired preening (Fig. 2G).

**Figure 1.**
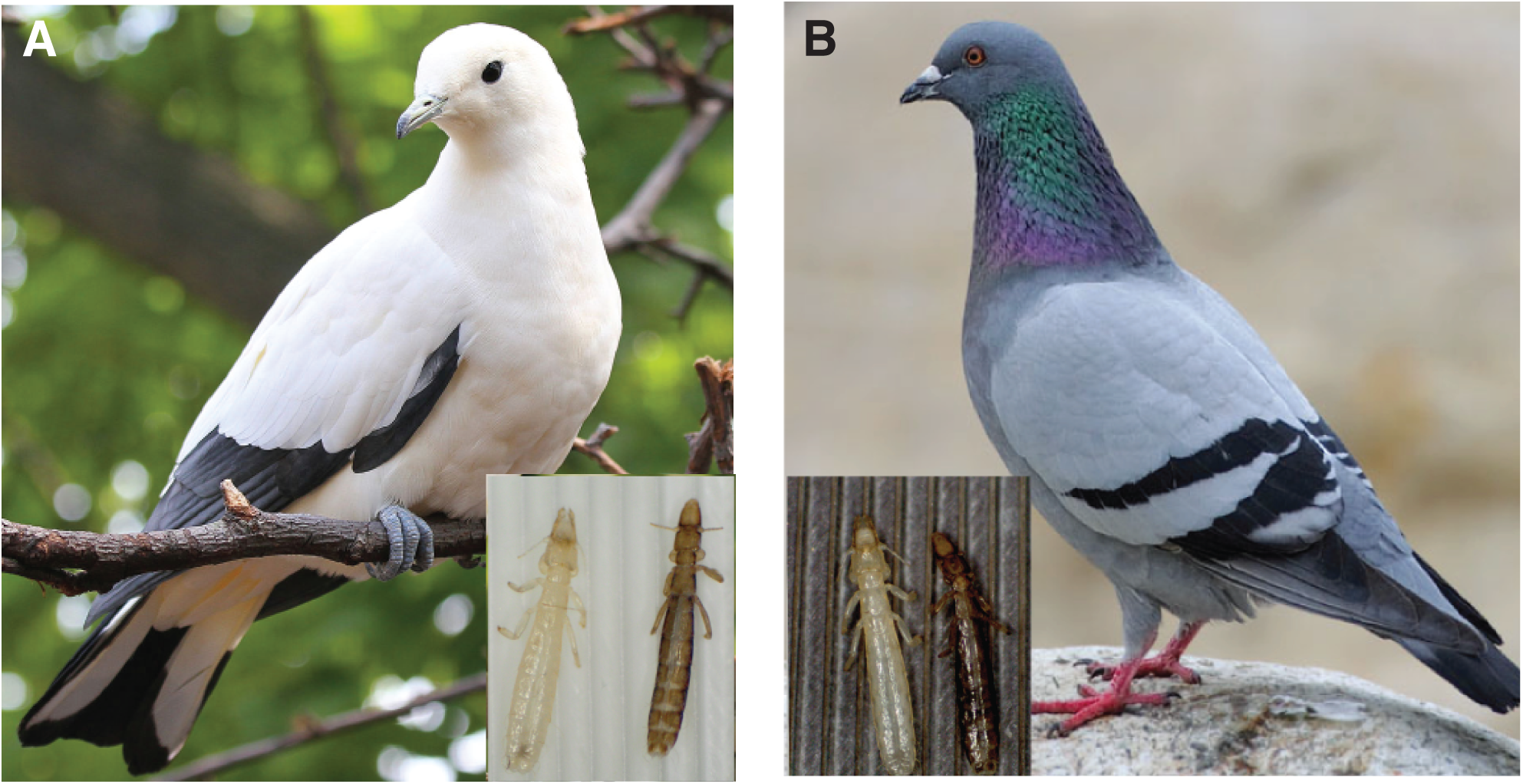
Matching coloration of host-specific feather lice on pigeons. The light-colored louse, *Columbicola wolffhuegeli* (left in both insets), parasitizes (A) the Australian pied imperial pigeon, *Ducula bicolor.* The dark-colored louse, *C. columbae* (right in both insets), parasitizes (B) the cosmopolitan rock pigeon, *Columba livia*. Pigeon photo (A) by Greg Hume (wikimedia.com); pigeon photo (B) by Mike Atkinson (mikeatkinson.net). Photos of lice by SEB.

**Figure 2.**
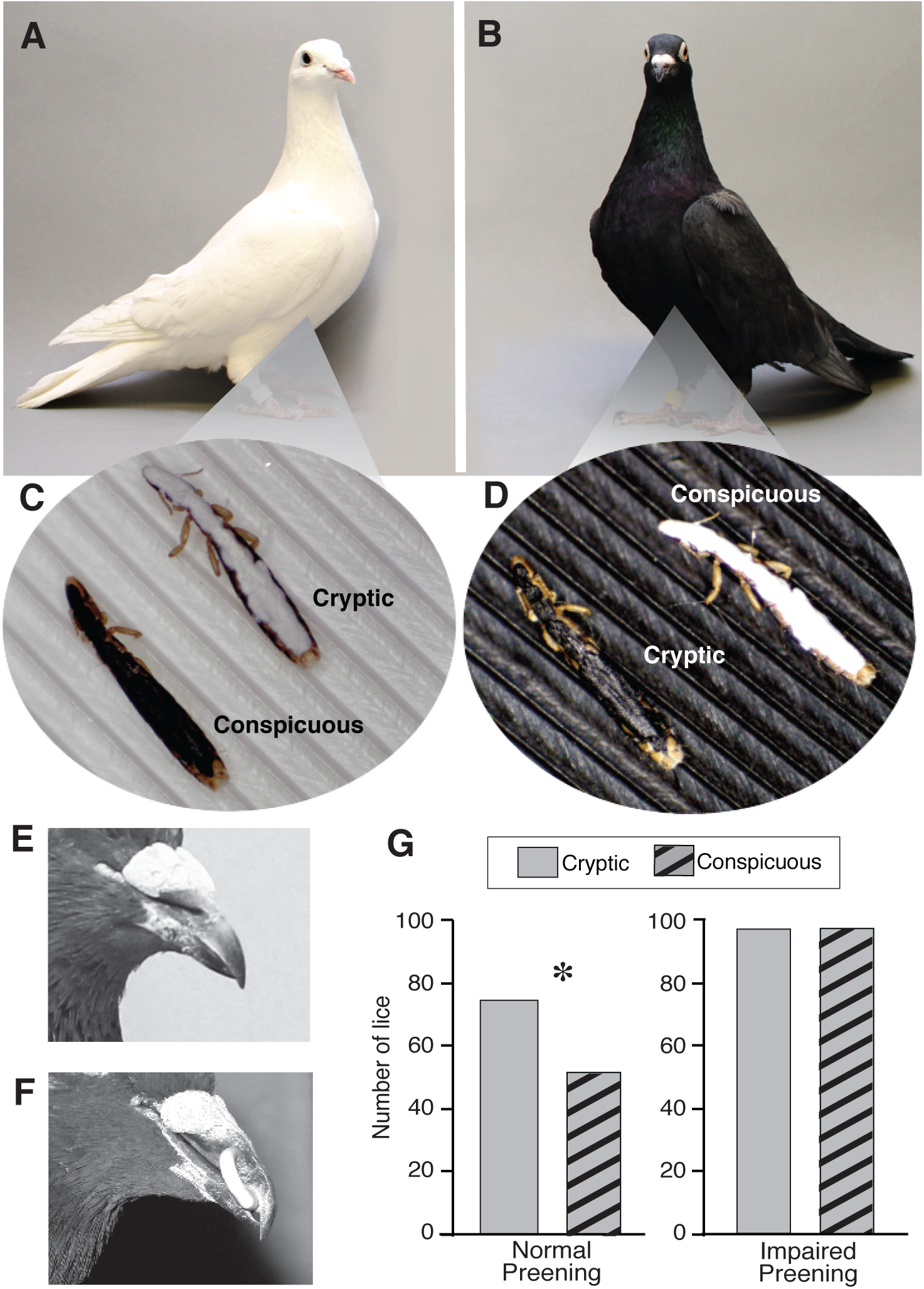
Preening selects for cryptically colored lice. Eight white (A) and eight black (B) rock pigeons were infested with live *C. columbae* that had been painted black or white to make them relatively conspicuous or cryptic, depending on background feather color (C, D). Each of the pigeons, which were isolated in 16 cages, received 30 conspicuous and 30 cryptic lice, for a total of 960 painted lice across the 16 birds. Half of the white pigeons and half of the black pigeons, chosen at random, could preen normally (E), while the other half were fitted with “poultry bits” to impair preening ability (F). (G) Pigeons that could preen normally had significantly more cryptic than conspicuous lice at the end of the 48-hr experiment (Fisher’s Exact test, P = 0.037). In contrast, there was no significant difference in the number of conspicuous and cryptic lice on pigeons with impaired preening (Fisher’s Exact test, P = 1.0).

This direct demonstration of preening-mediated selection for crypsis implies that preening leads to the diversification of parasite color among different colored hosts. Because feather lice are permanent parasites that pass their entire life cycle on the body of the host, they can be evolved experimentally under natural conditions on captive birds. Therefore, to test for adaptive divergence in response to host preening, we conducted a four-year experiment (ca. 60 louse generations) using *C. columbae* isolated on captive rock pigeons of different colors (see Materials and Methods). We transferred lice from wild caught grey feral rock pigeons (Fig. 1B) to white, black, or grey (control) rock pigeons that could either preen normally, or were impaired with bits.

At six-month intervals, random samples of lice were removed from each pigeon and digitally photographed under identical lighting conditions against a color standard [24]. The photographs were used to quantify the luminosity (brightness) of individual lice on a grey-scale from pixel values of 0 (pure black) to 255 (pure white) [25]. We used luminosity - the achromatic component of color - because feather lice vary mainly from light to dark [22]. We scored lice under visible light because they reflect little under UV light [22]. We also quantified variation in background coloration by measuring the luminosity of plumage on the white, black, and grey pigeons.

Over the course of the four-year experiment, the luminosity of lice on white and black birds changed relative to the luminosity of lice on control grey birds. The relative luminosity of lice on white birds increased dramatically, while the luminosity of lice on black birds decreased, but more slowly (Fig. 3A; Tables S1,S3). In contrast, lice on white and black birds with impaired preening showed no significant change in luminosity, relative to lice on control grey birds, even after 60 generations (Fig. 3B; Tables S2,S4). Thus, merely living and feeding on different colored feathers, in the absence of preening, had no effect on the color of the lice. Changes in the luminosity of lice on preening birds were proportional to differences in background luminosity, i.e. the luminosity of host plumage. The luminosity difference between grey and white plumage was five fold greater than that between grey and black plumage (Fig. 3C). Thus, lice on white birds presumably experienced more intense selection for background matching than lice on black birds. Differences in selection intensity may therefore have contributed to the greater change in the color of lice on white birds than on black birds.

**Figure 3.**
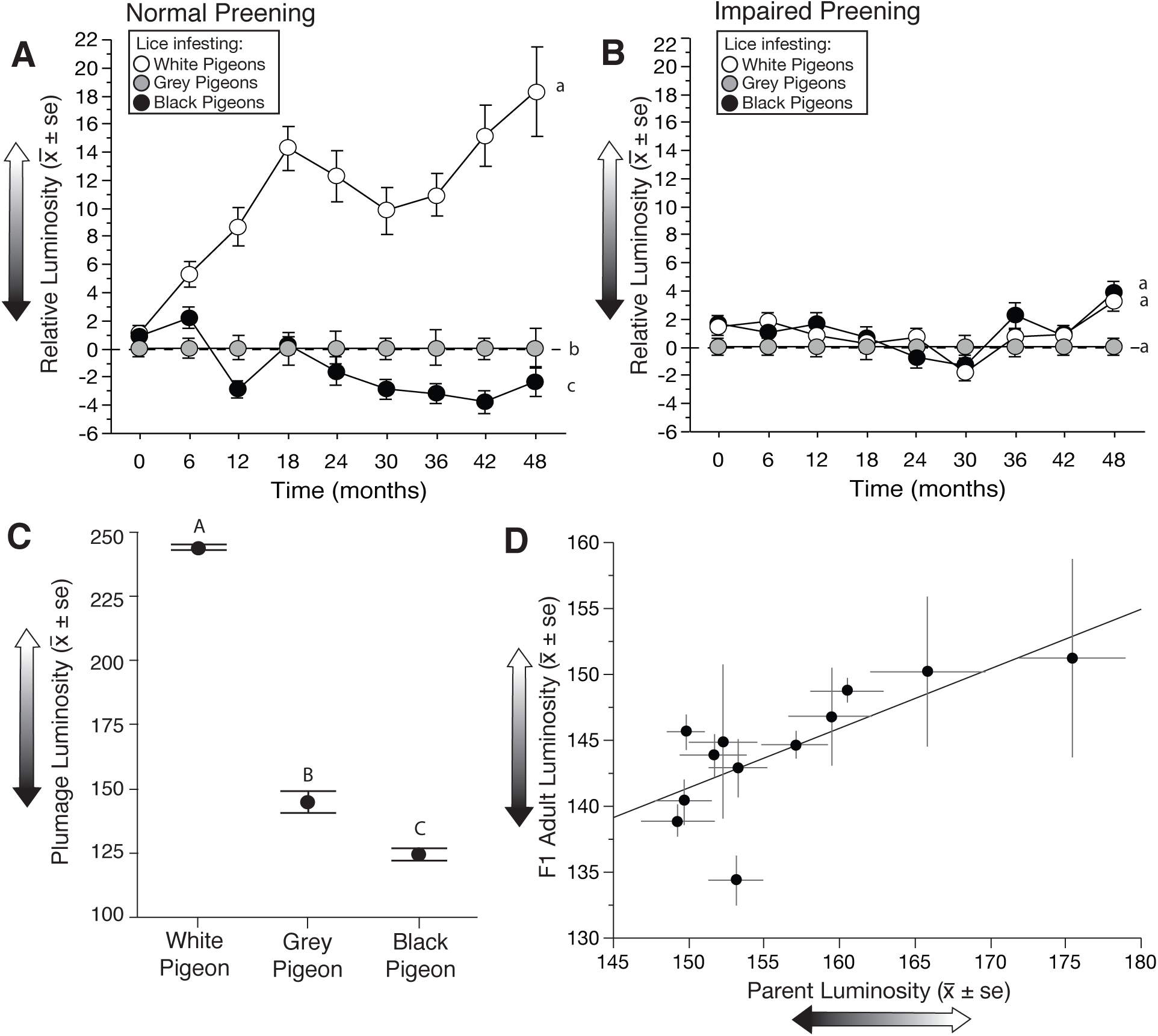
Evolution of feather lice *(C. columbae)* on different colored rock pigeons over a four-year period (ca. 60 louse generations). (A-B) The y-axis shows changes in the mean (± se) luminosity (brightness) of lice on white and black rock pigeons, relative to lice on grey rock pigeon controls (set to zero). Different lower-case letters indicate statistically significant differences. (A) On birds with normal preening, the relative luminosity of lice on white pigeons increased rapidly (LMM, P < 0.0001; Tables S1,S3); the relative luminosity of lice on black pigeons decreased, but more slowly (LMM, P = 0.001; Tables S1,S3). (B) Relative luminosity did not significantly change over time on white or black pigeons with impaired preening (LMM, P ≥ 0.34 in both cases, Tables S2,S4). (C) Luminosity of plumage from 5 white, 10 grey (control), and 8 black rock pigeons. The three groups differed significantly in luminosity (ANOVA df = 2,22, F = 242.6, P < 0.0001; Tukey-Kramer post hoc tests P < 0.001 for all possible comparisons). (D) Common garden experiment showing heritability of preening-induced changes in the color of lice evolved on different colored pigeons. The adult luminosity of parental and offspring cohorts of lice were highly correlated (linear regression: r = 0.72, df = 11, F = 11.7, P = 0.007).

To further investigate the basis of the observed color changes of lice on birds that could preen normally, we conducted a common garden experiment to test for heritability of the preening-mediated changes in color (see Materials and Methods). Two years into the four-year experiment, we removed random samples of lice from white, grey, and black pigeons, marked them by clipping several large hair-like setae, then transferred the lice to 12 bitted grey pigeons with impaired preening (common garden conditions for lice). Lice remained on the common garden birds for 48 days, which was sufficient time for the marked cohort of lice to breed and for their F1 lice to mature to the adult stage. All adult lice were then removed from each bird and digitally photographed to compare the color of the parental cohort (clipped lice) to adults of the offspring cohort (unclipped lice). Adult luminosity of the parental and offspring cohorts was highly correlated among the 12 common garden birds (Fig. 3D), thus demonstrating that color has a heritable component. This common garden experiment, together with the four-year experiment (Fig. 3 A,B), confirms that the lice on different colored birds evolved changes in luminosity in response to preening-mediated selection.

Over the four years of experimental evolution, the increase in mean luminosity of lice on normally preening white pigeons was also accompanied by an increase in variation (Fig. 4). By the end of the experiment, the range in luminosity of lice on white pigeons was much greater than that of lice on grey pigeons (Welch’s test of unequal variance *F* = 28.0, *df* = 1, 66.5, *P <*0.0001). This increase was generated by expansion at the upper end of the scale: 19 of the 50 lice on white pigeons (38.0%) were lighter than any of the lice on grey pigeons. The range in luminosity of experimentally evolved *C. columbae* on white pigeons overlapped the luminosity of *C. wolffhuegeli* (Figs. 4G), the host-specific parasite of the light-colored pied imperial pigeon.Remarkably, within 60 generations, lice evolved differences in color similar to those of related species diversified over millions of years. Thus, our study demonstrates rapid adaptive diversification in descendants of a single population living under natural conditions.

**Figure 4.**
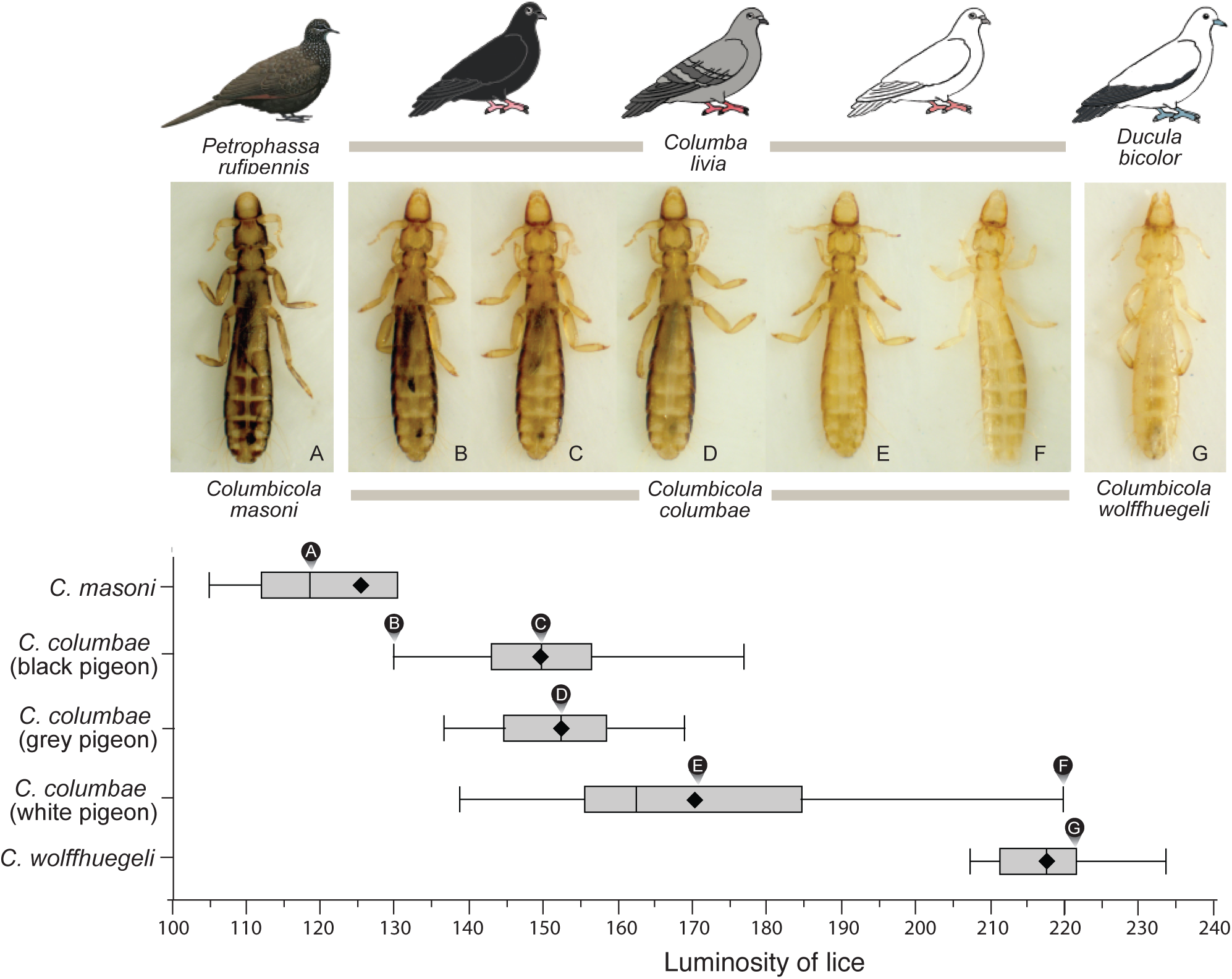
Luminosity of *C. columbae* after four years of experimental evolution on rock pigeons, compared to (A) *C. masoni* from a dark colored host species, the Australian chestnut-quilled pigeon (Petrophassa rufipennis), and (G) *C. wolffhuegeli* from a light colored host species, the Australian pied imperial pigeon (also shown in Fig. 1A). Images of experimentally evolved lice in B - F are: (B) one of the darkest C. *columbae* from a black rock pigeon, (C) *average* C. *columbae* from a black rock pigeon,(D) *average C. columbae* from a grey rock pigeon, (E) *average* C. columbae from a white rock pigeon, and (F) one of the lightest C. columbae from a white rock pigeon. Whisker plots show means (diamonds), medians (vertical lines), 1st and 3rd quartiles (boxes) and 1.5 interquartile ranges (whiskers). Letters above whisker plots indicate the luminosity of corresponding photographs. Sample sizes: 7 *C. masoni*, 18 C. wolffhuegeli, and 74, 36 and 50 C. *columbae* from black, grey and white rock pigeons, respectively. Photos of lice by JCA and SMV.

In summary, we show that preening selects for cryptic coloration, and causes the rapid divergence of heritable phenotypes on different host backgrounds. Thus, even small populations harbor sufficient additive genetic variation for rapid adaptation to novel hosts within several parasite generations. This process is integral to the successful establishment of parasite populations after rare episodes of dispersal to the “wrong” host species, the precursor to host switching. Adaptive radiation catalyzed by host switching is thought to be a central mechanism of diversification among parasites, which represent a substantial fraction of the earth’s biodiversity [6-8, 19]. Our results imply that other modes of host defense, such as immunological resistance, or secondary chemical compounds, may trigger rapid divergence in endoparasites, phytophagous insects, and other hyper-diverse groups. Host defense should be included with competition and predation as one of the principal mechanisms driving divergence in adaptive radiations.

## METHODS

Detailed methods are provided at the end of this manuscript.

## SUPPLEMENTAL INFORMATION

Supplemental information includes four tables and can be found at the end of this manuscript.

## ACKNOWLEDGEMENTS

We thank F. Adler, A. Beach, J. Baldwin-Brown, W. C. Brown, H. Campbell, K. Caseii, J. Endler, M. Evans, D. Feener, D. Kim, L. Mulvey, N. Phadnis, E. Poole, M. Reed, J. Ruff, J. Seger, N. Vickers, V. Zafferese and E. Waight for discussion and other assistance. All procedures followed guidelines of the Institutional Animal Care and Use Committee of the University of Utah. This work was supported by National Science Foundation DEB-0107947 and DEB-1342600.

## AUTHOR CONTRIBUTIONS

S.E.B., S.M.V., K.P.J., M.D.S., and D.H.C. designed the experiments. S.E.B., S.M.V. and J.C.A. collected the experimental data. S.E.B. and D.H.C. collected field specimens.S.E.B., S.M.V. and D.H.C. analyzed the data. S.E.B., K.P.J., M.D.S., and D.H.C. obtained funding for the work. S.E.B., S.M.V., K.P.J., M.D.S., and D.H.C. contributed to writing of the manuscript. S.E.B., S.M.V. and J.C.A. took photographs. S.E.B., S.M.V. and D.H.C. prepared figures.

## METHODS

### Study system

One of the challenges of experimental work with host-specific parasites is that, by definition, they are difficult to culture in sufficient numbers on novel host species. We circumvented this problem by working with rock pigeons (*Columba livia*), a single host species that harbors extensive intraspecific diversity in color as a result of artificial selection [26]. Rock pigeons with contrasting plumage colors were used as stepping stones to simulate environments that lice dispersing between different species of pigeons would encounter in nature. All animal procedures were approved by the IACUC of the University of Utah.

### Elimination of “background” lice

Before using pigeons in experiments, all “background” lice were eradicated by housing the birds in low humidity conditions (< 25% relative ambient humidity) for ≥ 10 weeks. This method kills lice and their eggs, while avoiding residues from insecticides [27]. During experiments, the relative humidity in animal rooms was increased to 35-60%, which provides sufficient humidity for feather lice to extract the moisture they need from the air while living on birds [28].

### Impaired preening

Preening was impaired using harmless poultry bits, which are C-shaped pieces of plastic inserted between the upper and lower mandibles of a bird’s beak (Fig. 2F). Bits spring shut in the nostrils to prevent dislodging, but without damaging the tissue. They create a 1 - 3 mm gap that prevents the forceps-like action of the bill required for efficient preening [29]. Bits have no apparent side effects and they do not impair the ability of birds to feed [30].

### Preening-mediated selection experiment

#### Painting lice

We manipulated the color of lice by covering the dorsal surface of live, adult *C. columbae* with enamel paint (Floquil™ Railroad Enamel, Vernon Hills, IL). Painting is a reliable method that has been used successfully with other species of lice, even under field conditions [31]. First, we tested whether the paint affects survival of *C. columbae*, as follows: live adult *C. columbae* were removed from infested pigeons and placed in a petri dish next to a small piece of dry ice to keep them anesthetized with CO_2_ during the painting process. We divided 75 lice into three treatments: 25 lice painted white, 25 lice painted black, and 25 unpainted control lice. Paint was applied to the dorsal surface of each louse with a very fine brush. Lice in the control treatment were handled and brushed, but without applying paint. We did not paint the legs of lice, as this might interfere with mobility. Painted lice were placed on feathers from grey feral pigeons in 50ml glass tubes in a Percival© incubator set at optimal conditions for lice: 33°C and 75% relative humidity on a 12-hour light/dark photoperiod [32]. We compared the survival of lice among the three treatments under a dissecting scope (Olympus SZ-CTV stereoscope) on six occasions over a 20-day period (days: 1, 3, 5, 7, 11, and 20). Painting had no significant effect on the survival of lice over this period of time (Kaplan-Meier Survival, Wilcoxon ?^2^=2.2, df = 2, *P* = 0.34).

#### Experimental infestation with painted lice

16 pigeons (half white, half black) were randomly divided (within each color treatment) into two preening treatments: half could preen normally, and the other half had their preening impaired with bits (see above). Pigeons in this experiment were housed individually in 30×30×56cm wire mesh cages in our animal facility. Cages were separated by plastic partitions to prevent any contact between the feathers of birds in adjacent cages, which might allow transmission of lice between pigeons. Birds were maintained on a 12-hour light/dark photoperiod and provided *ad libitum* grain, grit, and water.

Each bird received 30 cryptic lice and 30 conspicuous lice. The survival of lice was assessed 48 hours after the pigeons were experimentally infested. To do this, all pigeons were sacrificed and their lice removed by “body washing” [23]. Each louse was inspected under a dissecting scope and the number of white and black lice recovered from each bird was recorded.

### Experimental evolution

To test whether preening selects for divergence in the color of lice, we infested different colored pigeons with normal, unpainted *C. columbae*. Prior to experimental infestation, recipient pigeons were cleared of lice by housing them in low humidity conditions (as described above). Next, we transferred 2,400 lice from wild caught feral rock pigeons to 96 captive rock pigeons (25 lice per bird): 32 white pigeons, 32 black pigeons, and 32 grey pigeons (controls). Within each color, half the birds, chosen at random, were allowed to preen normally (Fig. 2E), whereas the other half were given bits to impair their preening (Fig. 2F). At this time (Time 0), we also randomly sampled hundreds of lice from the source population on wild caught grey feral pigeons and their luminosity was scored (as described below).

Pigeons were housed in groups of four in 1.8 × 1.5 × 1.0m aviaries. In summary, the 96 pigeons used in this experiment were housed in 24 aviaries, each containing four birds of the same color and preening treatment (two males and two females per aviary).

During the experiment, all pigeons were maintained on a 12-hour light/dark photoperiod and provided *ad libitum* grain, grit, and water. When a bird died during the course of the experiment (a rare occurrence), the lice from the dead bird were transferred to a new parasite-free pigeon of the same color and sex within 24 hours. *Columbicola columbae* lice can survive for days on a dead bird, but cannot leave the dead bird’s feathers under their own power, so few lice were lost.

The experiment ran for four years. Given that *C. columbae* has a mean generation time of 24.4 days [27], this is about 60 generations. Every six months, random samples of lice were removed from pigeons and digitally photographed. Lice were removed by anesthetizing them with CO_2_ [33]. After exposure to CO_2_ the feathers of each bird were ruffled over a collection tray. The lice were then photographed by placing each louse dorsal side up on a glass slide fitted with a Kodak® Q-13 white color standard. The lice were harmlessly immobilized by placing a 22×22mm micro cover slip (VWR®) directly on their body. Digital photographs were taken at high resolution (uncompressed TIFF 2560×1920 pixels) using a DP25 digital color camera on an Olympus SZ-CTV stereoscope linked to a computer running CellSens® image acquisition and analysis software. All of the photos were scored digitally [24, 25]. For each image, the metathorax was selected and luminosity calculated using the open source imaging software ImageJ 1.3. To correct for slight differences in luminosity due to variation in ambient lighting, we also recorded the luminosity of the color standard immediately adjacent to each louse. We determined how much the photograph of the color standard differed from pure white (luminosity = 255), then added this correction factor to the luminosity score [22].

### Plumage coloration

We also used digital photography to quantify the plumage luminosity of 23 dead pigeons representing the three colors in the evolution experiment (8 black pigeons, 5 white pigeons, and 10 grey pigeons). The dorsal abdomen and back of each bird was photographed next to a Kodak Q-13® white color standard. High-resolution (5184 x 3456 pixels) digital photos were taken using a Canon EOS Rebel SL1® digital camera. We then highlighted the plumage in each image with the “Quick Selection Tool” in Adobe Photoshop CC 2015®, and determined the luminosity of the highlighted area. To correct for slight differences in luminosity due to variation in ambient lighting, we also recorded the luminosity of the color standard immediately adjacent to the bird. We determined how much the photograph of the color standard differed from pure white (luminosity = 255), then added this correction factor to the luminosity score for the plumage [22]. We used the mean plumage luminosity of the dorsal and ventral surfaces of each pigeon in analyses.

### Common garden experiment

The goal of this experiment was to test for heritability of preening-mediated changes in the color of lice. We therefore used lice only from the 12 aviaries containing birds with normal preening. Lice of each sex were randomly sampled from each aviary using CO_2_. We marked the lice by clipping setae along the right side of the abdomen and thorax with scissors designed for retinal surgery. Setal clipping is a reliable method that has been used successfully with other species of lice, including under field conditions [34]. Removal of setae does not influence survival, and the setae do not grow back. After clipping, lice from each aviary were placed on a single grey pigeon with impaired preening. The 12 “common garden” pigeons were isolated in 12 wire mesh cages (30×30×56cm). After a period of 48 days, all lice were removed from each of the 12 pigeons using CO_2_ (as described above). At 48 days, most F1 offspring had developed to the adult stage, and could be distinguished from members of the parental cohort, which had clipped setae. In contrast, F2 lice had not yet developed to the adult stage. Thus, using this design, we were able to compare adults of the parental and F1 cohorts of lice on each common garden bird. F1 lice were removed from each bird and digitally photographed and their luminosity scored (as described above). The luminosity of the F1 cohort from each common garden pigeon (n = 4-29 lice per bird for a total of 170 lice) was then compared to the luminosity of the parental cohort (n = 11-48 lice per aviary, for a total of 303 lice).

## SUPPLEMENTAL INFORMATION

**Table S1:** Linear mixed model (LMM) summary comparing the luminosity of lice on pigeons with ***normal preening***. This LMM is based on the luminosity measurements of 3423 lice sampled over the four-year experiment. Luminosity data at Time 0 are from a random sample of lice drawn from the starting population. Luminosity data for the rest of the experiment (Time 6 mo. - Time 48 mo.) are for lice sampled from 48 individual birds (16 grey control pigeons, 16 white pigeons, and 16 black pigeons) housed in 4 aviaries (4 birds per aviary) for each color treatment. The intercept of the model is set to the value of grey control pigeons at the beginning of the experiment.

| ***Effect of host color on louse luminosity*** |  |  |  |  |  |
| --- | --- | --- | --- | --- | --- |
| <u>Random effects</u> | <u>Variance</u> | <u>Std. Dev.</u> |  |  |  |
| Individual Bird | 3.182 | 1.784 |  |  |  |
| Aviary | 3.560 | 1.887 |  |  |  |
| <u>Fixed effects</u> | <u>Estimate</u> | <u>Std. Err.</u> | <u>df</u> | <u>t value</u> | <u>Pr(>\|t\|)</u> |
| Intercept | 147.332 | 1.127 | 9.141 | 130.736 | < 0.001* |
| Breed (Black Pigeon) | -0.253 | 1.584 | 8.926 | -0.160 | 0.877 |
| Breed (White Pigeon) | 7.480 | 1.586 | 8.970 | 4.715 | 0.001* |

| ***Effect of host color on louse luminosity over time*** |  |  |  |  |  |
| --- | --- | --- | --- | --- | --- |
| <u>Random effects</u> | <u>Variance</u> | <u>Std. Dev.</u> |  |  |  |
| Individual Bird | 1.950 | 1.397 |  |  |  |
| Aviary | 1.786 | 1.336 |  |  |  |
| <u>Fixed effects</u> | <u>Estimate</u> | <u>Std. Err.</u> | <u>df</u> | <u>t value</u> | <u>Pr(>\|t\|)</u> |
| Intercept | 143.969 | 0.923 | 13.000 | 156.000 | < 0.001* |
| Time | 0.228 | 0.024 | 3354.000 | 9.334 | < 0.001* |
| Time x Breed (Black Pigeon) | -0.107 | 0.033 | 3354.000 | -3.249 | 0.001* |
| Time x Breed (White Pigeon) | 0.286 | 0.033 | 3356.000 | 8.592 | < 0.001* |

\* Indicates significance

**Table S2:** Linear mixed model (LMM) summary comparing the luminosity of lice on pigeons with ***impaired preening***. This LMM is based on the luminosity measurements of 5377 lice sampled over the four-year experiment (Table S4). Luminosity data at Time 0 are from a random sample of lice drawn from the starting population. Luminosity data for the rest of the experiment (Time 6 mo. - Time 48 mo.) are for lice sampled from 48 individual birds (16 grey control pigeons, 16 white pigeons, and 16 black pigeons) housed in 4 aviaries (4 birds per aviary) for each color treatment. The intercept of the model is set to the value of lice on grey control pigeons at the beginning of the experiment.

| ***Effect of host color on louse luminosity*** |  |  |  |  |  |
| --- | --- | --- | --- | --- | --- |
| <u>Random effects</u> | <u>Variance</u> | <u>Std. Dev.</u> |  |  |  |
| Individual Bird | 0.052 | 0.229 |  |  |  |
| Aviary | 0.533 | 0.730 |  |  |  |
| <u>Fixed effects</u> | <u>Estimate</u> | <u>Std. Err.</u> | <u>df</u> | <u>t value</u> | <u>Pr(>\|t\|)</u> |
| Intercept | 148.542 | 0.435 | 9.055 | 341.623 | < 0.001* |
| Breed (Black Pigeon) | 1.033 | 0.617 | 9.151 | 1.676 | 0.128 |
| Breed (White Pigeon) | 0.895 | 0.613 | 8.967 | 1.459 | 0.179 |

| ***Effect of host color on louse luminosity over time*** |  |  |  |  |  |
| --- | --- | --- | --- | --- | --- |
| <u>Random effects</u> | <u>Variance</u> | <u>Std. Dev.</u> |  |  |  |
| Individual Bird | < 0.001 | < 0.001 |  |  |  |
| Aviary | 0.602 | 0.776 |  |  |  |
| <u>Fixed effects</u> | <u>Estimate</u> | <u>Std. Err.</u> | <u>df</u> | <u>t value</u> | <u>Pr(>\|t\|)</u> |
| Intercept | 144.700 | 0.534 | 19.000 | 270.876 | < 0.001* |
| Time | 0.173 | 0.013 | 5364.000 | 13.046 | < 0.001* |
| Time x Breed (Black Pigeon) | 0.018 | 0.019 | 5363.000 | 0.952 | 0.341 |
| Time x Breed (White Pigeon) | 0.003 | 0.019 | 5367.000 | 0.173 | 0.863 |

\* Indicates significance

**Table S3:**
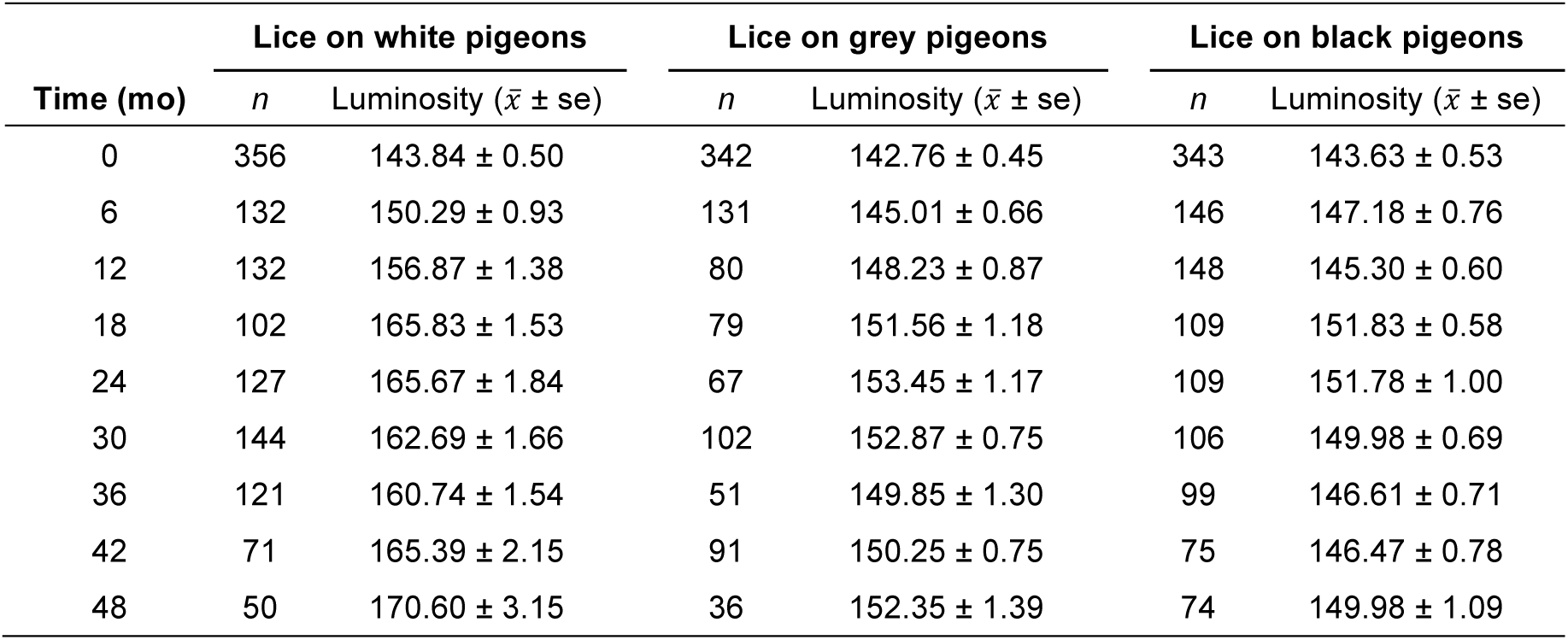
Mean luminosity of lice from white, grey, and black pigeons with ***normal preening*** over the course of the four-year experiment. Luminosity data at Time 0 are from a random sample of lice drawn from the starting population. Luminosity data for the rest of the experiment (Time 6 mo. - Time 36 mo.) are for lice sampled from 48 individual birds (16 grey control pigeons, 16 white pigeons, and 16 black pigeons) housed in 4 aviaries (4 birds per aviary) for each color treatment.

**Table S4:** Mean luminosity of lice from black, white, and grey pigeons with ***impaired preening*** over the course the four-year experiment. Luminosity data at Time 0 data are from a random sample of lice drawn from the starting population. Luminosity data for the rest of the experiment (Time 6 mo. - Time 36 mo.) are from lice sampled from 48 individual birds (16 grey control pigeons, 16 white pigeons, and 16 black pigeons) housed in 4 aviaries (4 birds per aviary) for each color treatment.

| Time (mo) | Lice on white pigeons |  | Lice on grey pigeons |  | Lice on black pigeons |  |
| --- | --- | --- | --- | --- | --- | --- |
| | *n* | Luminosity ($\bar{x} \pm se$) | *n* | Luminosity ($\bar{x} \pm se$) | *n* | Luminosity ($\bar{x} \pm se$) |
| 0 | 348 | 143.86 ± 0.45 | 332 | 142.44 ± 0.46 | 351 | 144.09 ± 0.56 |
| 6 | 194 | 146.42 ± 0.52 | 186 | 144.60 ± 0.55 | 185 | 145.61 ± 0.70 |
| 12 | 192 | 149.98 ± 0.63 | 186 | 149.18 ± 0.69 | 160 | 150.80 ± 0.86 |
| 18 | 186 | 152.12 ± 0.63 | 162 | 151.90 ± 0.84 | 164 | 152.56 ± 0.82 |
| 24 | 195 | 150.53 ± 0.55 | 183 | 149.87 ± 0.64 | 161 | 149.31 ± 0.74 |
| 30 | 193 | 151.10 ± 0.51 | 166 | 152.92 ± 0.81 | 173 | 151.57 ± 0.76 |
| 36 | 195 | 150.27 ± 0.57 | 193 | 149.56 ± 0.65 | 171 | 151.81 ± 0.92 |
| 42 | 193 | 151.26 ± 0.51 | 189 | 150.39 ± 0.59 | 184 | 151.17 ± 0.68 |
| 48 | 162 | 154.75 ± 0.65 | 195 | 151.53 ± 0.57 | 178 | 155.38 ± 0.79 |

## REFERENCES

1. Simpson, G. G. (1953). The Major Features of Evolution (Columbia University Press).

2. Schluter, D. (2000). The Ecology of Adaptive Radiation (Oxford University Press).

3. Losos, J. B. (2010). Adaptive radiation, ecological opportunity, and evolutionary determinism. American Naturalist 175, 623–639.

4. Nosil, P., & Crespi, B. J. (2006). Experimental evidence that predation promotes divergence in adaptive radiation. Proceedings of the National Academy of Sciences of the United States of America 103, 9090–9095.

5. Meyer, J. R., & Kassen, R. (2007). The effects of competition and predation on diversification in a model adaptive radiation. Nature 446, 432–435.

6. Price, P. W. (1980). Evolutionary Biology of Parasites (Princeton University Press).

7. de Meeus, T., & Renaud, F. (2002). Parasites within the new phylogeny of eukaryotes. Trends in Parasitology 18, 247–251.

8. Poulin, R. (2014). Parasite biodiversity revisited: frontiers and constraints. International Journal for Parasitology 44, 581–589.

9. Wiens, J. J., Lapoint, R. T., & Whiteman, N. K. (2015). Herbivory increases diversification across insect clades. Nature Communications 6, 8370.

10. Jezkova, T., & Wiens, J. J. (2017). What explains patterns of diversification and richness among animal phyla? American Naturalist 189, 201–212.

11. Clayton, D. H., Bush, S. E., & Johnson, K. P. (2015). Coevolution of Life on Hosts: Integrating Ecology and History (University of Chicago Press).

12. Nylin, S., Agosta, S., Bensch, S., Boeger, W. A., Braga, M. P., Brooks, D. R., et al. (2017). Embracing colonizations: a new paradigm for species association dynamics. Trends in Ecology and Evolution 33, 4–14.

13. Giraud, T., Gladieux, P., & Gavrilets, S. (2010). Linking the emergence of fungal plant diseases with ecological speciation. Trends in Ecology and Evolution 25, 387–395.

14. Johnson, K. P., Weckstein, J. D., Meyer, M. J., & Clayton, D. H. (2011). There and back again: switching between host orders by avian body lice (Ischnocera: Goniodidae). Biological Journal of the Linnean Society 102, 614–625.

15. Fordyce, J. A. (2010). Host shifts and evolutionary radiations of butterflies. Proceedings of the Royal Society B 277, 3735–3743.

16. Hardy, N. B., & Otto, S. P. (2014). Specialization and generalization in the diversification of phytophagous insects: tests of the musical chairs and oscillation hypotheses. Proceedings of the Royal Society B 281, 20132960.

17. Ehrlich, P. R., & Raven, P. H. (1964). Butterflies and plants: a study in coevolution. Evolution 18, 586–608.

18. Zietara, M. S., & Lumme, J. (2002). Speciation by host switch and adaptive radiation in a fish parasite genus Gyrodactylus (Monogenea, Gyrodactylidae). Evolution 56, 2445–2458.

19. Forbes, A. A., Devine, S. N., Hippee, A. C., Tvedte, E. S., Ward, A. K. G., Widmayer, H. A., & Wilson, C. J. (2017). Revisiting the particular role of host shifts in initiating insect speciation. Evolution 71, 1126–1137.

20. Loker, E. S. (2012). Macroevolutionary immunology: a role for immunity in the diversification of animal life. Frontiers in Immunology 3, 25–46.

21. Harbison, C. W., & Clayton, D. H. (2011). Community interactions govern host-switching with implications for host–parasite coevolutionary history. Proceedings of the National Academy of Sciences 108, 9525.

22. Bush, S. E., Kim, D., Reed, M., & Clayton, D. H. (2010). Evolution of cryptic coloration in ectoparasites. American Naturalist 176, 529–535.

23. Clayton, D. H., & Drown, D. M. (2001). Critical evaluation of five methods for quantifying chewing lice (Insecta: Phthiraptera). Journal of Parasitology 87, 1291–1300.

24. Villafuerte, R., & Negro, J. (1998). Digital imaging for colour measurement in ecological research. Ecology Letters 1, 151–154.

25. Stevens, M., Párraga, C. A., Cuthill, I. C., Partridge, J. C., & Troscianko, T. S. (2007). Using digital photography to study animal coloration. Biological Journal of the Linnean Society 90, 211–237.

26. Shapiro, M. D., & Domyan, E. T. (2013). Domestic pigeons. Current Biology 23, R302–R303.

27. Harbison, C. W., Bush, S. E., Malenke, J. R., & Clayton, D. H. (2008). Comparative transmission dynamics of competing parasite species. Ecology 89, 3186–3194.

28. Nelson, B. C., & Murray, M. D. (1971). The distribution of Mallophaga on the domestic pigeon (Columba livia). International Journal for Parasitology 1, 21–29.

29. Clayton, D. H., Moyer, B. R., Bush, S. E., Jones, T. G., Gardiner, D. W., Rhodes, B. B., & Goller, F. (2005). Adaptive significance of avian beak morphology for ectoparasite control. Proceedings of the Royal Society B: Biological Sciences 272, 811.

30. Clayton, D. H., & Tompkins, D. M. (1995). Comparative effects of mites and lice on the reproductive success of rock doves (Columba livia). Parasitology 110, 195–206.

31. Zohdy, S., Kemp, A. D., Durden, L. A., Wright, P. C., & Jernvall, J. (2012). Mapping the social network: tracking lice in a wild primate (Microcebus rufus) population to infer social contacts and vector potential. BMC Ecology 12, 4.

32. Bush, S. E., & Clayton, D. H. (2006). The role of body size in host specificity: reciprocal transfer experiments with feather lice. Evolution 60, 2158–2167.

33. Moyer, B. R., Gardiner, D. W., & Clayton, D. H. (2002). Impact of feather molt on ectoparasites: looks can be deceiving. Oecologia 131, 203–210.

34. Durden, L. A. (1983). Sucking louse (Hoplopleura erratica: Insecta, Anoplura) exchange between individuals of a wild population of eastern chipmunks, Tamias striatus, in central Tennessee, USA. Journal of Zoology, London 201, 117–123.

